## Supplemental Material for "Automating clinical assessments of memory deficits: Deep Learning based scoring of the Rey-Osterrieth Complex Figure"

### Supplementary Material

#### Image resolution analysis

The ROCF images in our dataset have varying resolutions, ranging from 100 x 140 pixels to 3500 x 5300 pixels. Since our models are trained on a fixed input resolution, we investigated the effect of different resolutions on the model performance measured in terms of MAE. To that end, we trained the multilabel classification model with and without data augmentation for inputs of size 78x100, 116x150, 232x300 and 348x450. In principle, when using smaller images, then more information is lost due to resizing, while, on the other hand, a resolution which is too large requires bigger models due to increased complexity in the underlying distribution. In addition, using our data augmentation pipeline with small images might negatively affect the performance since the interpolation techniques used in the semantic transformations like rotations, potentially leads to an additional loss of information. We observed this in our experiments which showed that inputs of size 232x300 yielded the best performance, both for the model with and without data augmentation (Figure 2E). Thus, all subsequent analyses were performed with images of size 232x300.

| | MAE | MSE | $R^2$ |
|---|---|---|---|
| Classifier TTA | 1.259 [1.214, 1.308] | 3.902 [3.598, 4.268] | 0.963 |
| Classifier NA | 1.228 [1.185, 1.275] | 3.700 [3.418, 4.042] | 0.965 |
| Regressor DA | 1.162 [1.122, 1.204] | 3.174 [2.958, 3.425] | 0.970 |
| Classifier DA | 1.161 [1.118, 1.206] | 3.368 [3.105, 3.743] | 0.968 |
| Regressor DA + TTA | 1.152 [1.110, 1.194] | 3.152 [2.928, 3.399] | 0.970 |
| Classifier DA + TTA | 1.149 [1.106, 1.194] | 3.315 [3.061, 3.646] | 0.969 |
| Regressor TTA | 1.139 [1.099, 1.181] | 3.074 [2.850, 3.335] | 0.971 |
| Regressor NA | 1.138 [1.098, 1.180] | 3.078 [2.850, 3,354] | 0.971 |
| Final Model (Best Regressor + Best Classifier) | 1.113 [1.074, 1.156] | 3.000 [2.775, 3,262] | 0.972 |

Table S1. The performance metrics for all model variants. NA: non-augmented, DA: data augmentation is performed during training, TTA: test-time augmentation. The 95% confidence interval is shown in square brackets.

| | MAE | MSE | $R^2$ | | MAE | MSE | $R^2$ |
|---|---|---|---|---|---|---|---|
| **Item 1** | 0.104 | 0.095 | 0.800 | **Item 10** | 0.051 | 0.048 | 0.950 |
| **Item 2** | 0.170 | 0.166 | 0.587 | **Item 11** | 0.081 | 0.072 | 0.853 |
| **Item 3** | 0.148 | 0.151 | 0.778 | **Item 12** | 0.101 | 0.092 | 0.869 |
| **Item 4** | 0.127 | 0.132 | 0.730 | **Item 13** | 0.116 | 0.107 | 0.778 |
| **Item 5** | 0.125 | 0.132 | 0.744 | **Item 14** | 0.094 | 0.092 | 0.839 |
| **Item 6** | 0.148 | 0.131 | 0.802 | **Item 15** | 0.073 | 0.090 | 0.899 |
| **Item 7** | 0.063 | 0.073 | 0.923 | **Item 16** | 0.094 | 0.101 | 0.862 |
| **Item 8** | 0.105 | 0.090 | 0.860 | **Item 17** | 0.091 | 0.078 | 0.869 |
| **Item 9** | 0.091 | 0.084 | 0.893 | **Item 18** | 0.114 | 0.108 | 0.814 |
| | | | | **Total score** | 1.113 | 3.000 | 0.972 |

Table S2. Per-item and total performance estimates for the final model. MAE, MSE and $R^2$ are estimated directly from the estimated scores.

| Score Interval | MSE | MAE | Score Interval | MSE | MAE | Score Interval | MSE | MAE | Score Interval | MSE | MAE |
|---|---|---|---|---|---|---|---|---|---|---|---|
| (0 - 0.5) (n=22) | 3.72 | 1.52 | (10 - 10.5) (n=104) | 5.52 | 1.79 | (19 - 19.5) (n=42) | 3.07 | 1.25 | (28 - 28.5) (n=130) | 3.28 | 1.27 |
| (1 - 1.5) (n=15) | 2.78 | 1.17 | (11 - 11.5) (n=33) | 3.17 | 1.41 | (20 - 20.5) (n=131) | 4.90 | 1.56 | (29 - 29.5) (n=92) | 3.11 | 1.28 |
| (2 - 2.5) (n=52) | 3.15 | 1.41 | (12 - 12.5) (n=103) | 5.77 | 1.74 | (21 - 21.5) (n=45) | 4.41 | 1.30 | (30 - 30.5) (n=155) | 3.26 | 1.28 |
| (3 - 3.5) (n=26) | 3.93 | 1.52 | (13 - 13.5) (n=28) | 2.77 | 1.36 | (22 - 22.5) (n=105) | 3.75 | 1.44 | (31 - 31.5) (n=134) | 3.33 | 1.38 |
| (4 - 4.5) (n=62) | 3.66 | 1.52 | (14 - 14.5) (n=108) | 4.82 | 1.56 | (23 - 23.5) (n=75) | 3.26 | 1.05 | (32 - 32.5) (n=167) | 2.67 | 1.15 |
| (5 - 5.5) (n=32) | 7.23 | 2.09 | (15 - 15.5) (n=48) | 3.70 | 1.52 | (24 - 24.5) (n=120) | 3.43 | 1.25 | (33 - 33.5) (n=196) | 1.73 | 0.96 |
| (6 - 6.5) (n=92) | 5.27 | 1.61 | (16 - 16.5) (n=141) | 5.17 | 1.51 | (25 - 25.5) (n=75) | 2.45 | 1.03 | (34 - 34.5) (n=299) | 1.21 | 0.77 |
| (7 - 7.5) (n=21) | 6.96 | 1.74 | (17 - 17.5) (n=54) | 4.40 | 1.56 | (26 - 26.5) (n=149) | 4.32 | 1.44 | (35 - 35.5) (n=295) | 0.88 | 0.58 |
| (8 - 8.5) (n=88) | 4.79 | 1.52 | (18 - 18.5) (n=115) | 6.03 | 1.51 | (27 - 27.5) (n=93) | 2.82 | 1.11 | (36) (n=564) | 0.45 | 0.20 |
| (9 - 9.5) (n=34) | 2.21 | 1.13 | | | | | | | | | |

Table S3. Performance per total score interval. Thirty seven intervals were evaluated, across the whole range of scores. We evaluated MAE and MSE for the total score within each interval.

| | MAE | MSE | $R^2$ | | MAE | MSE | $R^2$ |
|---|---|---|---|---|---|---|---|
| **Item 1** | 0.118 | 0.108 | 0.800 | **Item 10** | 0.049 | 0.060 | 0.952 |
| **Item 2** | 0.103 | 0.122 | 0.588 | **Item 11** | 0.081 | 0.070 | 0.848 |
| **Item 3** | 0.132 | 0.126 | 0.810 | **Item 12** | 0.115 | 0.103 | 0.862 |
| **Item 4** | 0.082 | 0.111 | 0.758 | **Item 13** | 0.077 | 0.078 | 0.808 |
| **Item 5** | 0.084 | 0.121 | 0.761 | **Item 14** | 0.095 | 0.088 | 0.841 |
| **Item 6** | 0.166 | 0.142 | 0.805 | **Item 15** | 0.080 | 0.083 | 0.925 |
| **Item 7** | 0.070 | 0.083 | 0.932 | **Item 16** | 0.058 | 0.080 | 0.899 |
| **Item 8** | 0.103 | 0.091 | 0.878 | **Item 17** | 0.082 | 0.069 | 0.900 |
| **Item 9** | 0.077 | 0.074 | 0.917 | **Item 18** | 0.086 | 0.081 | 0.881 |
| | | | | **Total score** | 1.131 | 3.319 | 0.969 |

Table S4. Per-item and total performance estimates for the final model with prospective data. MAE, MSE and $R^2$ are estimated directly from the estimated scores.

| Score Interval | MSE | MAE | Score Interval | MSE | MAE | Score Interval | MSE | MAE | Score Interval | MSE | MAE |
|---|---|---|---|---|---|---|---|---|---|---|---|
| (0 - 0.5) (n=3) | 14.83 | 3.33 | (10 - 10.5) (n=35) | 5.81 | 1.84 | (19 - 19.5) (n=21) | 2.77 | 1.21 | (28 - 28.5) (n=138) | 3.96 | 1.36 |
| (1 - 1.5) (n=4) | 9.55 | 2.06 | (11 - 11.5) (n=23) | 3.96 | 1.49 | (20 - 20.5) (n=58) | 5.96 | 1.7 | (29 - 29.5) (n=61) | 4.45 | 1.54 |
| (2 - 2.5) (n=7) | 7.79 | 2.29 | (12 - 12.5) (n=39) | 6.2 | 2.03 | (21 - 21.5) (n=30) | 5.08 | 1.63 | (30 - 30.5) (n=136) | 3.28 | 1.31 |
| (3 - 3.5) (n=5) | 4.09 | 1.85 | (13 - 13.5) (n=16) | 5.09 | 1.5 | (22 - 22.5) (n=79) | 5.29 | 1.65 | (31 - 31.5) (n=53) | 3.45 | 1.46 |
| (4 - 4.5) (n=20) | 9.3 | 2.11 | (14 - 14.5) (n=34) | 6.74 | 1.74 | (23 - 23.5) (n=37) | 3.49 | 1.24 | (32 - 32.5) (n=133) | 3.14 | 1.33 |
| (5 - 5.5) (n=6) | 14.76 | 2.88 | (15 - 15.5) (n=22) | 5.66 | 1.68 | (24 - 24.5) (n=97) | 5.55 | 1.73 | (33 - 33.5) (n=93) | 2.27 | 1.13 |
| (6 - 6.5) (n=18) | 7.24 | 1.99 | (16 - 16.5) (n=41) | 5.77 | 1.65 | (25 - 25.5) (n=58) | 4.23 | 1.53 | (34 - 34.5) (n=243) | 1.75 | 1.02 |
| (7 - 7.5) (n=8) | 14.88 | 3.0 | (17 - 17.5) (n=14) | 8.1 | 2.07 | (26 - 26.5) (n=94) | 2.72 | 1.29 | (35 - 35.5) (n=204) | 1.85 | 0.67 |
| (8 - 8.5) (n=31) | 7.35 | 1.96 | (18 - 18.5) (n=50) | 3.32 | 1.45 | (27 - 27.5) (n=55) | 5.34 | 1.59 | (36) (n=519) | 1.03 | 0.19 |
| (9 - 9.5) (n=10) | 3.79 | 1.32 | | | | | | | | | |

Table S5. Performance per total score interval with prospective data. Thirty seven intervals were evaluated, across the whole range of scores. We evaluated MAE and MSE for the total score within each interval.

| | | |
|---|---|---|
| Unit Correct | Placed Properly | 2 |
| | Placed Poorly | 1 |
| Unit Distorted, Incomplete but Recognisable | Placed Properly | 1 |
| | Placed Poorly | 0.5 |
| Absent of Unrecognisable | | 0 |

Supplementary Figure S1: Original scoring system according to Osterrieth

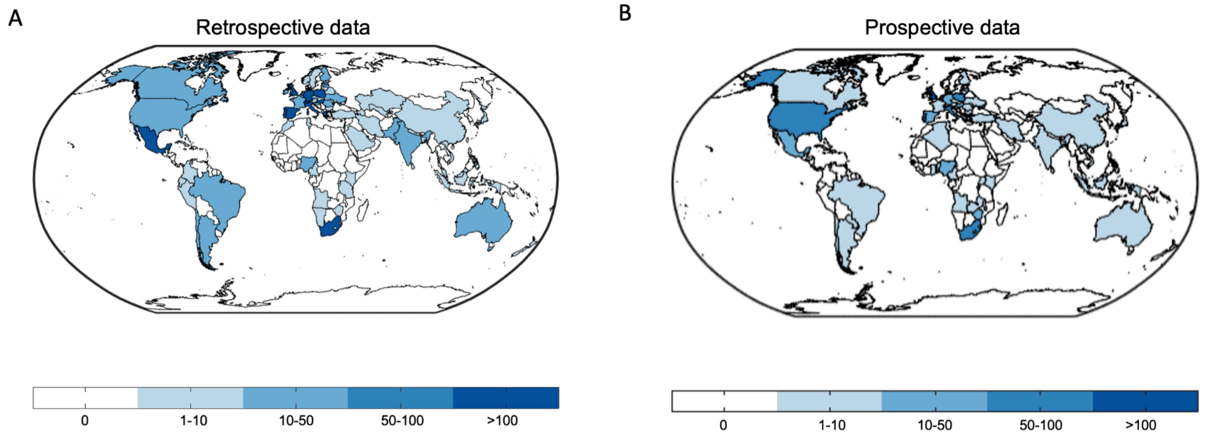


Supplementary Figure S2. World maps depict the worldwide distribution of the origin of the data. A: retrospective data B: prospective data.

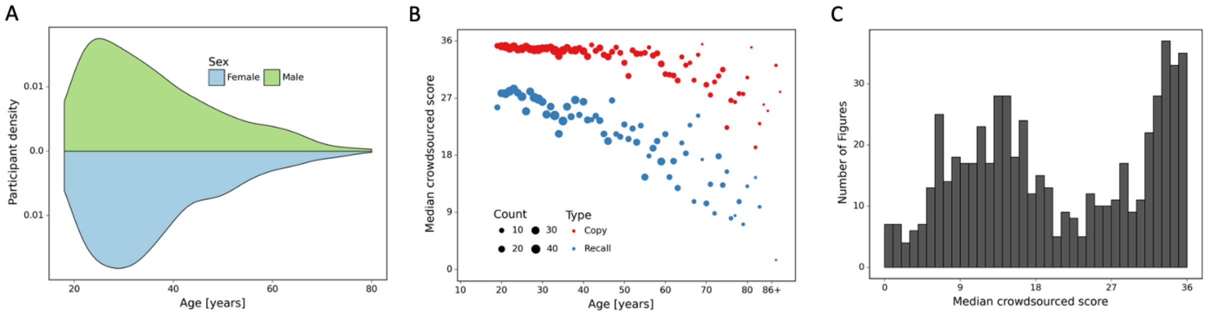


Supplementary Figure S3. A: demographics of the participants of the prospectively collected data. B: performance in the copy and (immediate) recall condition across the lifespan in the prospectively collected data. C: distribution of number of images for each total score for the prospectively collected data.

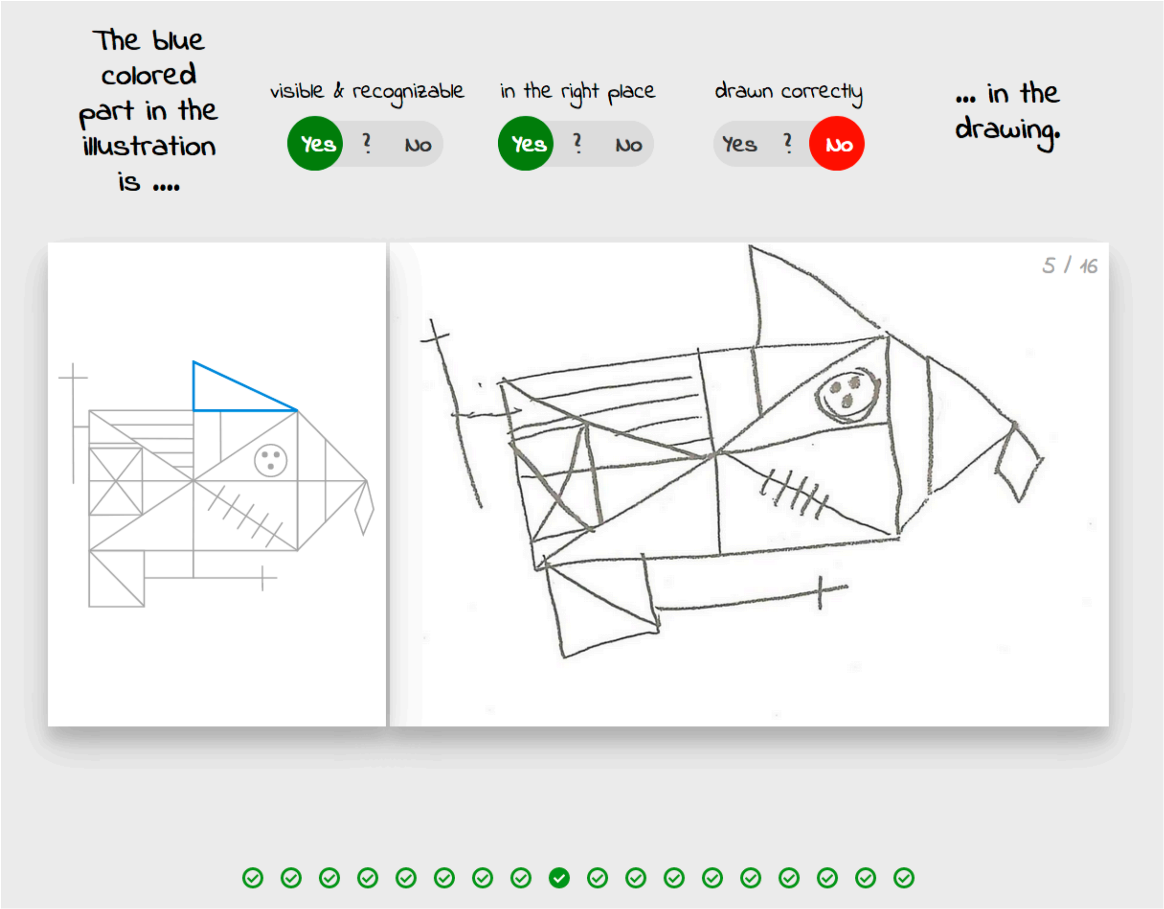


Supplementary Figure S4. The graphical user interface of the crowdsourcing application

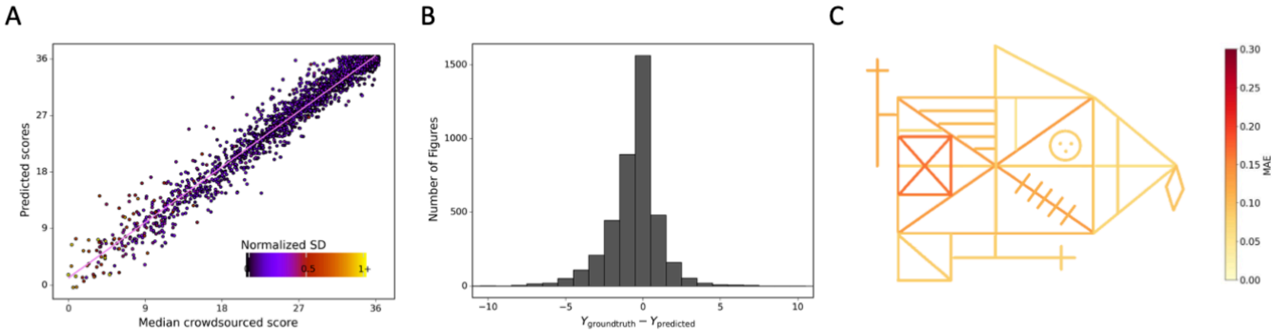


Supplementary Figure S5. Detailed performance of the model on the prospective data. Contrasting the ratings of our model. A: against the ground truth. A jitter is applied to better

highlight the dot density. B: The distribution of errors for our model on the prospective data are displayed. C: The MAE of our model is displayed for each individual item of the figure (see also supplementary table S4).

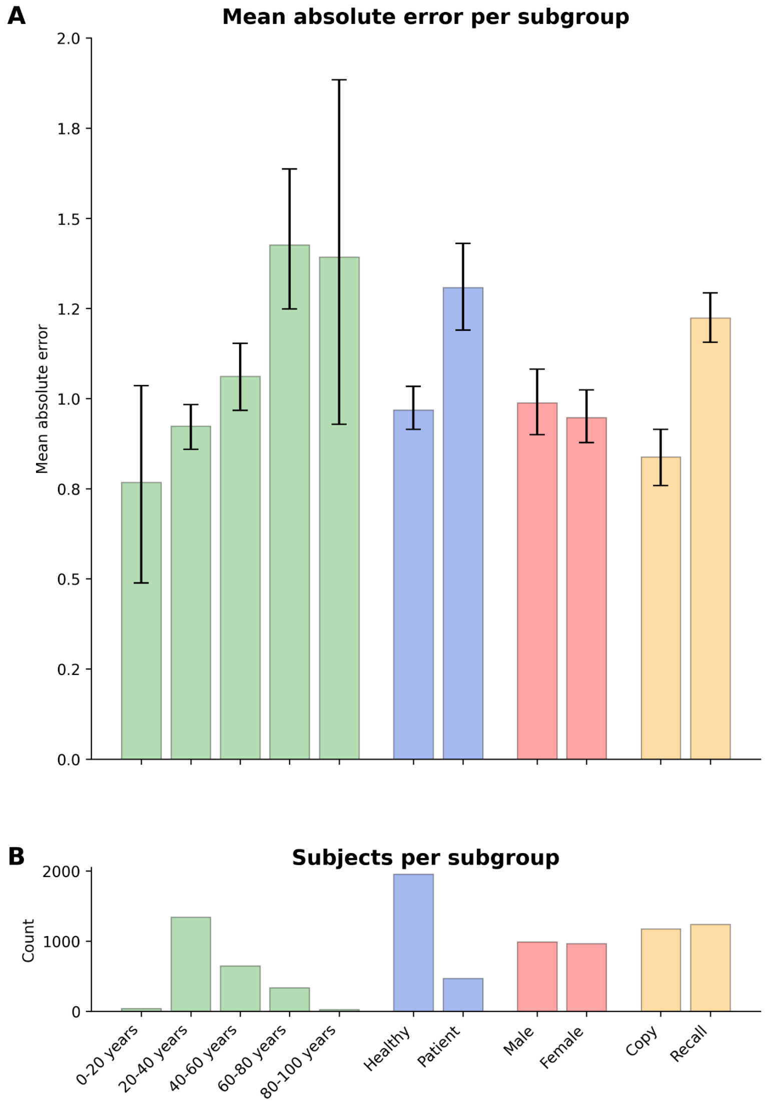


Supplementary Figure S6. A. Displayed are the mean absolute error and bootstrapped 95% confidence intervals of the model performance across different ROCF conditions (copy and recall), demographics (age, gender), and clinical statuses (healthy individuals and patients) for the prospective data. B. The number of subjects in each subgroup is depicted. Please note, that we did not have sufficient information on the specific patient diagnoses in the prospective data to decompose the model performance for specific clinical conditions.

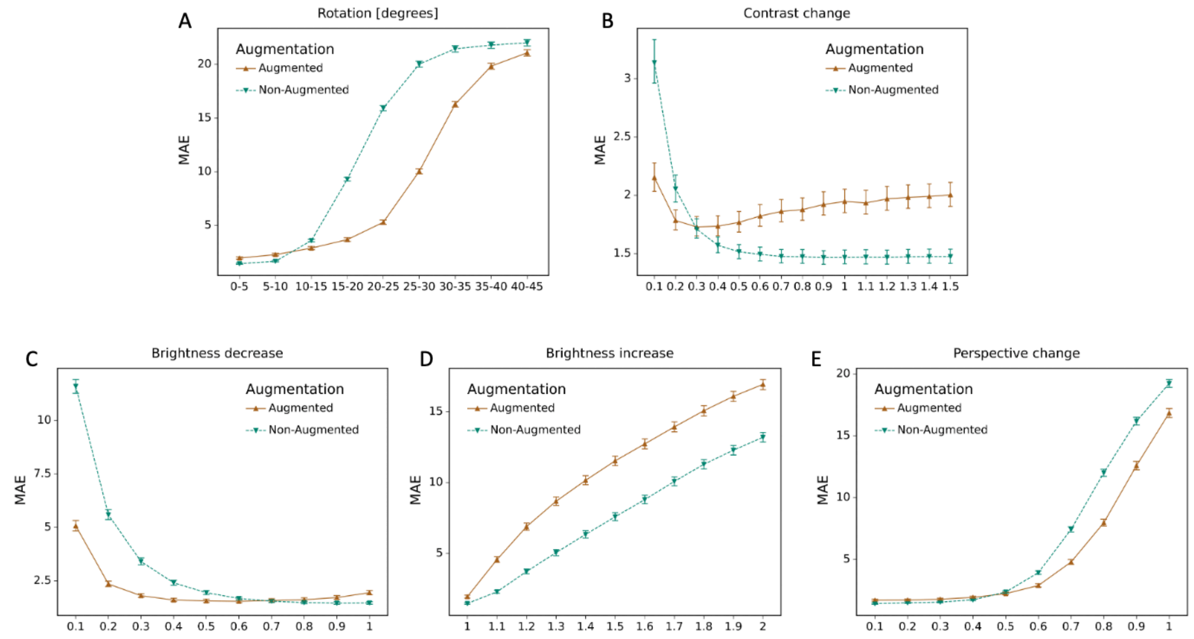


Supplementary Figure S7. Effect of data augmentation

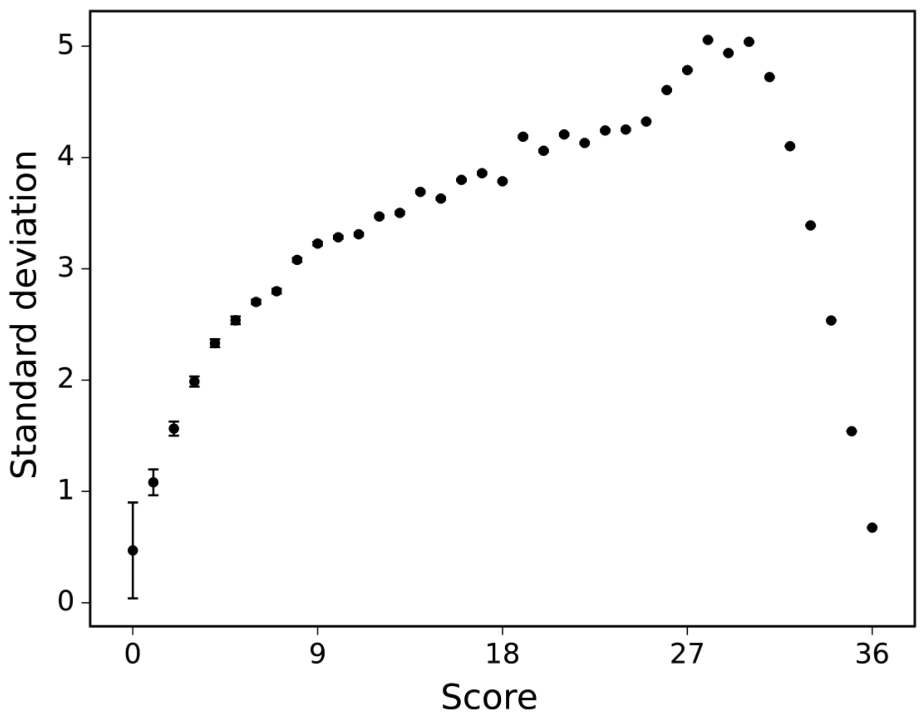


Supplementary Figure S8: The standard deviation of the human raters is displayed across differently scored drawings.

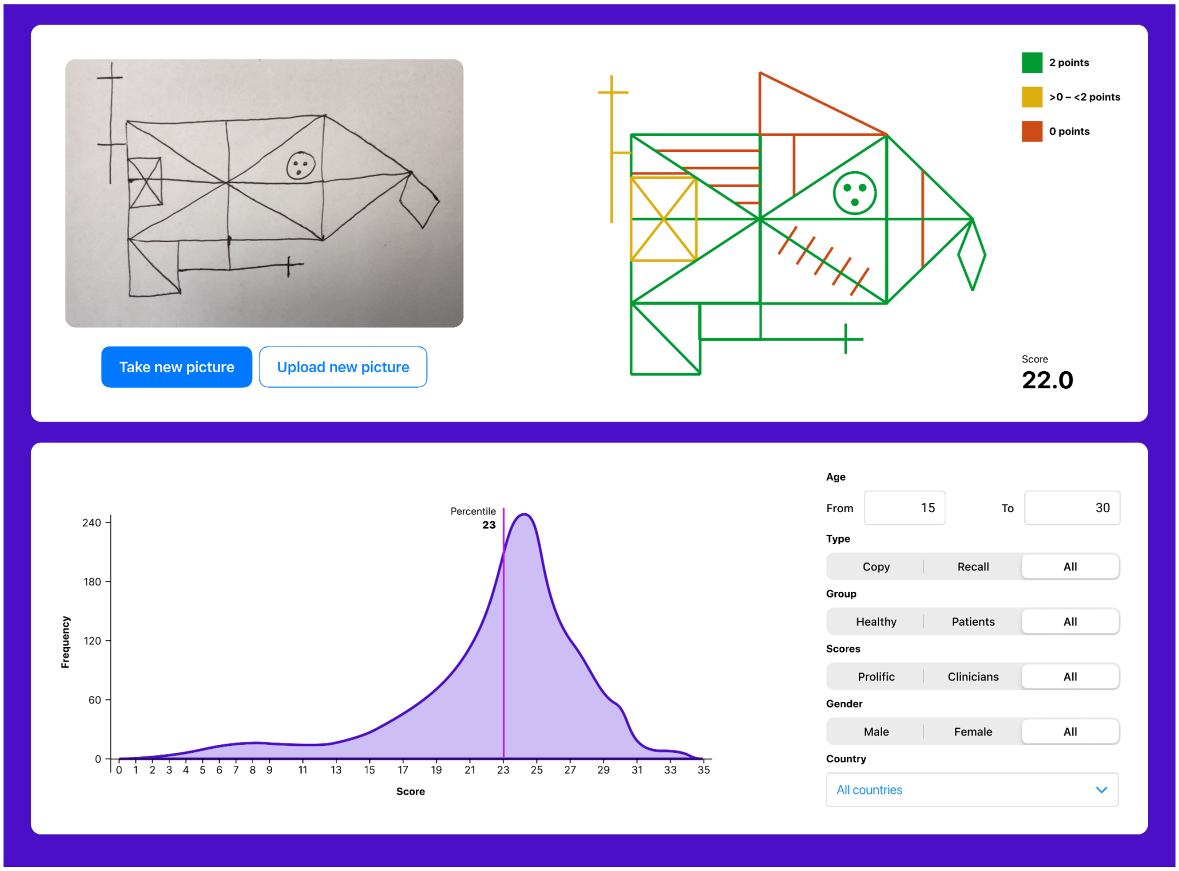


Supplementary Figure S9: The user interface for the tablet- (and smartphone-) based application. The application enables explainability by providing a score for each individual item. Furthermore, the total score is displayed. The user can also compare the individual with a choosable norm population.
